# Cuticular hydrocarbons are associated with mating success and insecticide resistance in malaria vectors

**DOI:** 10.1101/2021.04.20.440144

**Authors:** Kelsey L. Adams, Simon P. Sawadogo, Charles Nignan, Abdoulaye Niang, Douglas G. Paton, W. Robert Shaw, Adam South, Jennifer Wang, Maurice A. Itoe, Kristine Werling, Roch K. Dabiré, Abdoulaye Diabaté, Flaminia Catteruccia

## Abstract

*Anopheles coluzzii* females, important malaria vectors in Africa, mate only once in their lifetime. Mating occurs in aerial swarms with a high male-to-female ratio, where the traits underlying male mating success are largely unknown. Here, we investigated whether cuticular hydrocarbons (CHCs) influence mating success in natural mating swarms in Burkina Faso. As insecticides are widely used in this area for malaria control, we also determined whether CHCs affect insecticide resistance levels. We find that mated males have higher CHC abundance than unmated controls, suggesting CHCs could be a determinant of mating success. Additionally, mated males have higher insecticide resistance under pyrethroid challenge, and we show a link between resistance intensity and CHC abundance. Taken together, our results reveal overlapping roles played by CHCs in mate choice and insecticide resistance, and point to sexual selection for insecticide resistance traits that limit the efficacy of our best malaria control tools.

## Introduction

Two of the major malaria vectors in sub-Saharan Africa, *Anopheles gambiae* and *Anopheles coluzzii* of the *Anopheles gambiae* complex, are largely monandrous, which means the lifetime reproductive fitness of females depends on a single mating event^1^. In these anopheline species, mating occurs in aerial swarms where males heavily outnumber females, an indication of female-driven mate selection and/or scramble mating competition among males^2,3^. Swarm initiation remains poorly understood, but it is thought that these mosquitoes integrate visual signals from geographic markers and lighting, circadian cues, acoustic signals and volatile pheromones to identify the presence of conspecific individuals of the opposite sex^4–7^. When a swarming male approaches a female, there is substantial contact between their legs and abdomens after which, in successful mating events, the male grasps the female and completes copulation^8^. It has been demonstrated that during these close-range interactions, females can exhibit rejection behavior before copulation starts^4,9,10^. Leading up to these close range interactions, harmonic convergence, the adjustment of wing beat frequencies between a male and female, is observed in *Anopheles* like in other mosquito species, but there is no evidence that it increases the likelihood of successful copulation in *An. gambiae^11–13^*. Other studies have striven to understand whether male fitness, reflected by body size, determines mating outcomes, with unclear and conflicting conclusions^5,14–16^. So, in spite of substantial efforts, close-range cues involved in mate choice remain largely unknown in *Anopheles*.

Contact pheromones, including cuticular hydrocarbons (CHCs), are widely used by insects during social or sexual communication^17,18^. CHCs are waxy molecules derived from fatty acids via a biosynthetic process that involves desaturases, elongases, fatty acid synthases, and cytochrome P450 enzymes^*19,20*^. Biosynthesis occurs in specific cells called oenocytes, from where they are transported to the surface of the cuticle by lipophorin proteins, where they can regulate permeability in addition to playing pheromonal roles^21^. These compounds are chemically diverse, and are thought to be highly tuned to environmental pressures such as aridity as well as subject to sexual selection, resulting in plasticity in their composition and levels^22^. In mosquitoes, the role of CHCs in communication has not been fully elucidated, although reports indicate that stripping the cuticle with solvent^23–25^ or treating virgin females with CHC extracts from either males or females^26^ can reduce insemination rates, suggesting that CHCs may alter mosquito attractiveness. Further, recent data in *Anopheles stephensi* mosquitoes shows that males treated with the CHC heptacosane inseminate more females compared to untreated controls^6^, indicating a potential role for CHCs in mating success.

The possibility of sexual selection for CHCs in *Anopheles* is particularly interesting because it is already known that these traits are advantageous during selection by insecticide pressure. Cuticular insecticide resistance, a thickening of the cuticle caused by increased deposition of CHCs, cuticular proteins and chitin, leads to reduced or slowed insecticide penetrance^27–29^. Higher CHC levels in resistant mosquito populations have been linked to overexpression of two cytochrome P450 enzymes, *CYP4G16* and *CYP4G17*, that act as decarbonylases in the last steps in the CHC biosynthesis pathway^*27,28*^. Therefore, if CHCs are implicated in female mate choice or male competition during swarming, cuticular thickening due to selective pressures imposed by insecticides may also affect male mating success. Understanding whether CHCs affect both mating biology and insecticide resistance is a fascinating biological question intersecting evolution, ecology, and reproductive biology. This question is particularly relevant in areas of Africa where widespread insecticide resistance is threatening the efficacy of our best malaria control tools, which are predominantly based on the use of insecticides against vector species. Here we investigated whether CHCs are associated with male mating success and with insecticide resistance in field *An. coluzzii* populations from Burkina Faso. We show that males that successfully mate with females in natural mating swarms have higher total abundance of CHCs, and that these males survive longer during insecticide exposure. Moreover, we identify signatures of cuticular resistance in those populations and show their association with survival after insecticide exposure. Our data support a model by which CHCs play overlapping roles in mate choice and insecticide resistance and suggest that sexual selection for cuticular pheromone abundance may aid mosquitoes to withstand insecticide pressure. These findings have important repercussions for insecticide-based malaria control programs as well as for currently proposed genetic control strategies.

## Results

### Mated males have higher total CHC abundances in natural*An. coluzzii* swarms

To investigate whether CHC levels affect male mating success we decided to study males from natural mating swarms. This way we avoided using colonized mosquitoes which have adapted to confined laboratory conditions, a process likely to affect mating behavior and select for traits that are not necessarily relevant in field conditions^30^. To this end, we collected mated and unmated control mosquitoes from natural *An. coluzzii* swarms in VK7, a village in the Vallée du Kou area near Bobo Dioulasso, Burkina Faso. Mated males were collected *in copula*, while the control unmated groups were collected at different time points during the swarming period (throughout peak swarming, or at late time points) in random sweeps. Although we could not prove that control males had not mated during that evening, we expect this to be the case for two reasons: (1) return to the swarm on the same evening is unlikely, given the steep energy demands associated with copulation^14^, and (2) the highly biased sex ratios and large numbers of males in these swarms (several thousands) make sampling of returned males improbable. We therefore refer to these males as unmated controls, which are likely a reflection of the average swarming male.

We extracted CHCs from multiple pools of 5 males from either mated or unmated groups (**Figure 1A**), and extracts were submitted for gas chromatography mass spectrometry (GC-MS) analysis to retrieve quantitative and qualitative information of the CHC profiles. While all groups of males showed the same diversity of 38 CHC compounds regardless of mated status (**Supplementary Table 1**), mated males captured *in copula* had higher (by 1.37-fold) levels of CHCs compared to unmated controls from either peak or late time points after normalizing for wing length, a proxy for adult size **(Figure 1B).** No difference was instead detected between peak and late control groups (**Figure 1B**). When the two unmated groups were pooled, the greater CHC abundance of mated males was maintained **(Supplementary Figure 1A).** Wing length was not significantly different between mated and unmated groups in all comparisons **(Supplementary Figure 1B, C),** and we further showed that when incorporated into a Generalized Linear Model, mating group has a greater impact on CHC abundance compared to wing length **(Supplementary Table 2A)**. Among the 38 identified compounds, 15 had increased abundance in mated males compared to both peak and late unmated control groups (**Figure 1C**). The representation of each individual compound relative to the total abundance (proportional abundance) was similar between the three groups, indicating that the major differences between CHCs of these groups are quantitative and not qualitative **(Supplementary Figure 1D).** Although we cannot rule out the possibility that differences in age or life history could also be associated with mating success and CHC levels, these data suggest that CHC abundance may be a sexually selected trait.

**Figure 1.**
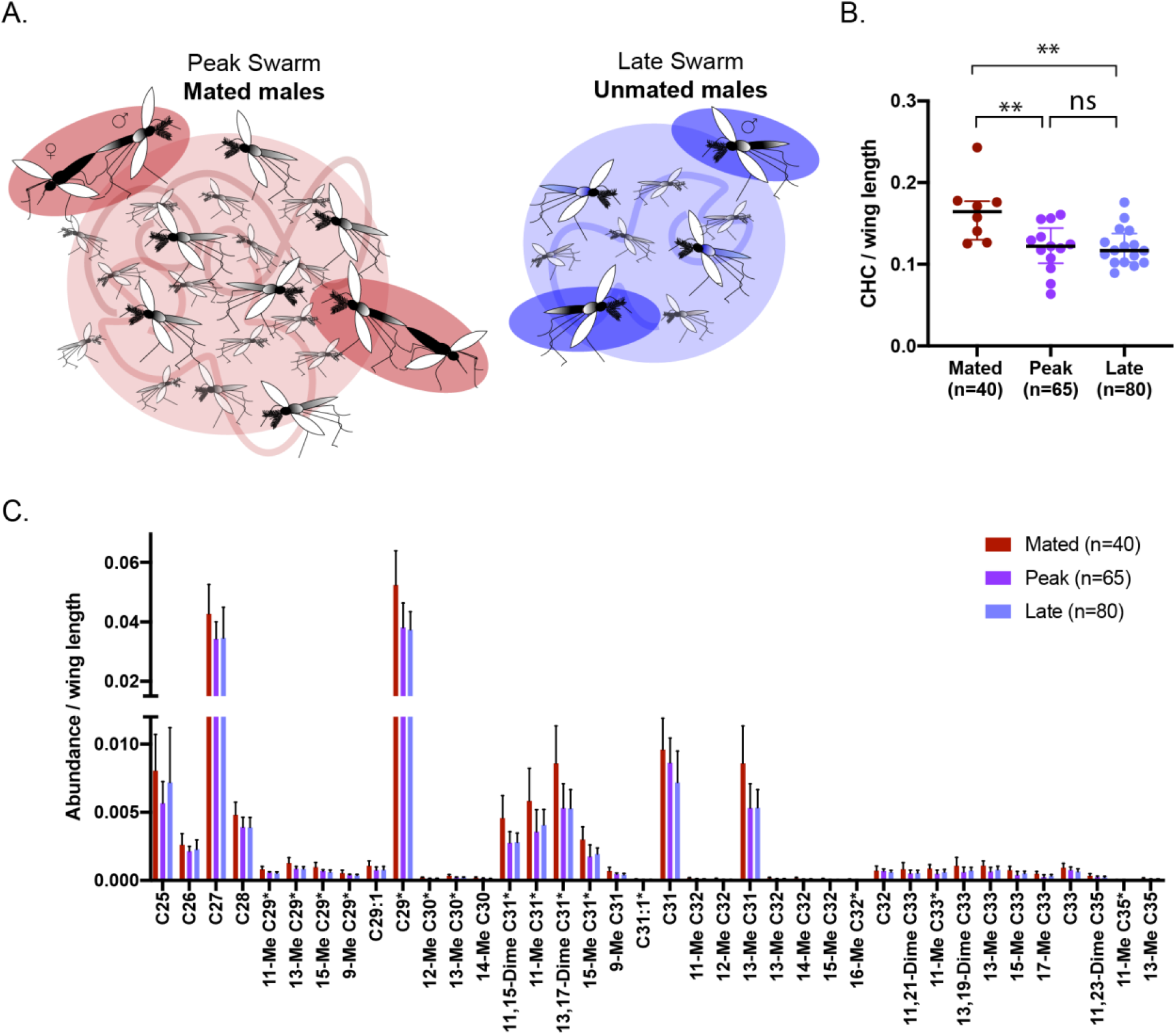
Successful *An. coluzzii* males in natural swarms have higher abundance of CHCs. (A) Scheme of captures of male groups from mating swarms, showing mated males (left image, red) and unmated controls collected late in the swarm (right image, blue). (B) Total abundance of CHCs is higher in mated compared to unmated males when captured at either peak or late time points during the swarm (Tukey’s multiple comparisons, *p*=0.0045 (mated vs peak unmated), *p*=0.0041 (mated vs late unmated)). The mean sum of response ratios for all CHCs divided by the mean wing length for each sample is shown. Error bars represent SD, and *n* describes total number of mosquitoes. (C) Mated males have higher abundance of 15 of 38 CHCs compared to both unmated males captured at the peak or late time point detected by GC-MS, shown here as the median response ratio to a pentadecane internal standard and normalized to wing length. Nomenclature for each compound indicates position of methyl (Me) or Dimethyl (Dime) groups on the carbon chain. Error bars represent interquartile ranges; Benjamini-Hochberg corrected *p* values from Mann-Whitney tests are displayed in full in Supplementary Table 1. Asterisks are indicated next to names of compounds with statistically significant differences in both peak and late unmated groups compared to the mated group.

### Mated males from natural mating swarms survive longer under permethrin exposure

Based on these findings, and evidence that CHCs are linked with insecticide resistance^27^, we next directly investigated whether mated males also have higher resistance to permethrin, an insecticide widely used on Long Lasting Insecticide-treated Nets (LLINs) in this area^31^. We collected mated and unmated (from the late time point) males from natural swarms as above **(Figure 1A)**. Given that no difference appears between CHCs of peak versus late unmated controls we decided to compare only the late group in the remainder of the study. Our reasoning is that the late time point is the most relevant control as by then no more females join the swarm and therefore males in this group are highly unlikely to mate that same evening. The day following collection we exposed these two groups to a 3.75% (5X) continuous dose of permethrin using WHO bioassay cylinders and permethrin-impregnated papers^32^. Mosquitoes were monitored for knockdown (failure to fly) every 30 minutes, and their total time to knockdown (referred to here as “survival”) was recorded **(Figure 2A upper panel).** When exposed to this dose of permethrin, mated males were 1.65 times more likely to survive the exposure (log-rank test, *p*=0.036), with a median survival 30 minutes longer than unmated controls **(Figure 2B).**

We reasoned that a lower permethrin dose may give us higher resolution on the differences between these mosquitoes, and so we next utilized an intermittent exposure regime using a 1.875% (2.5X) dose of permethrin, and also included recovery periods between exposures **(Figure 2A lower panel)**. When exposed to this dose, mated males were again more likely to survive (by 3.72 times) compared to unmated controls (log rank test, *p*<0.0001) **(Figure 2C)**. While control males had a median time to death of 90 minutes, in the mated group more than half of the males were still alive after 210 minutes of exposure, so their median time to death was undefined. Although we again observed no significant difference in wing length of mated compared to unmated males **(Supplementary Figure 2)**, we nonetheless incorporated wing length into a proportional hazards model and determined that this was not a significant factor in predicting survival time in Figure 2B **(Supplementary Table 2B)**. Wing lengths were not paired to survival time in males from Figure 2C, so this analysis could not be performed. When combined, our results show that males that are successful at mating exhibit greater resistance to insecticides.

**Figure 2.**
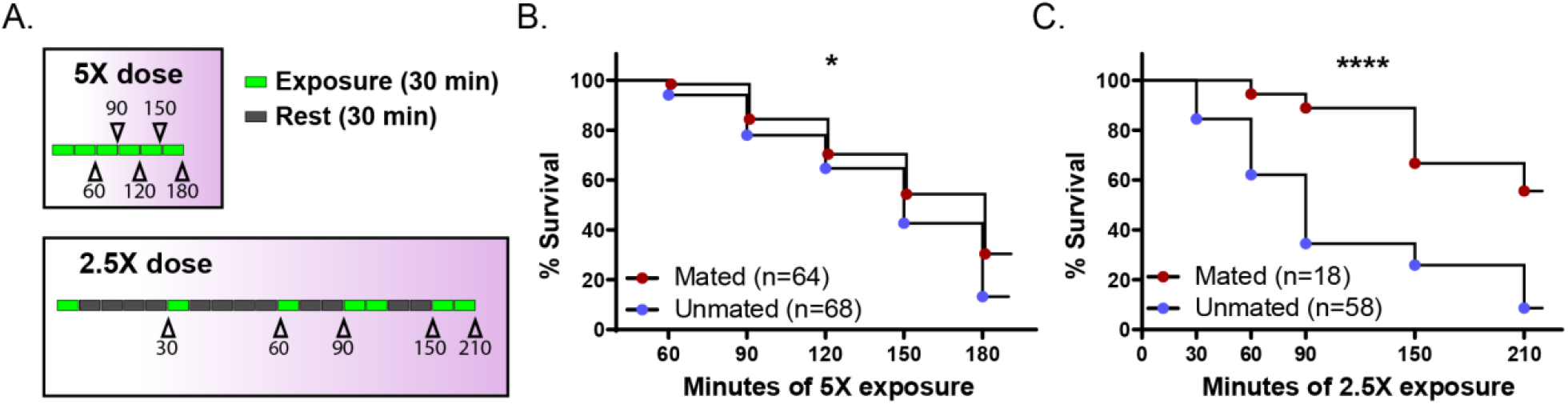
Successful males have higher permethrin resistance. Mated and control unmated males were captured from natural swarms as in Figure 1A. (A) These males were exposed to either a continuous 5X dose of permethrin (upper panel) or a series of 30 or 60 minute 2.5X permethrin exposures (lower panel) and monitored for knockdown after each exposure. Schematics represent 30-minute time intervals as either exposure periods (green) or rest periods (grey). Arrowheads denote time points for survival monitoring, labeled according to the cumulative minutes of permethrin exposure. Mated males survived longer on average to these permethrin exposures compared to unmated males for both the (B) 5X (*p*=0.036) and (C) 2.5X (*p*<0.0001) doses (log-rank tests). n represents total number of mosquitoes.

### *An. coluzzii* populations from Vallée du Kou show evidence of cuticular resistance

The evidence of a relationship between CHC abundance, mating success and resistance to pyrethroids prompted us to determine whether cuticular resistance is a mechanism acting in these mosquito populations, as suggested by other studies^28,33–35^. We first confirmed high intensity of pyrethroid resistance in this region by exposing adult *An. coluzzii* mosquitoes to permethrin, after collecting them as larvae from natural breeding sites in the Vallée du Kou. These tests detected nearly 100% survival at the standard 0.75% (1X) permethrin dose, and >70% survival at both 2.5X and 5X doses **(Figure 3A)**, in agreement with previous reports^31^.

**Figure 3.**
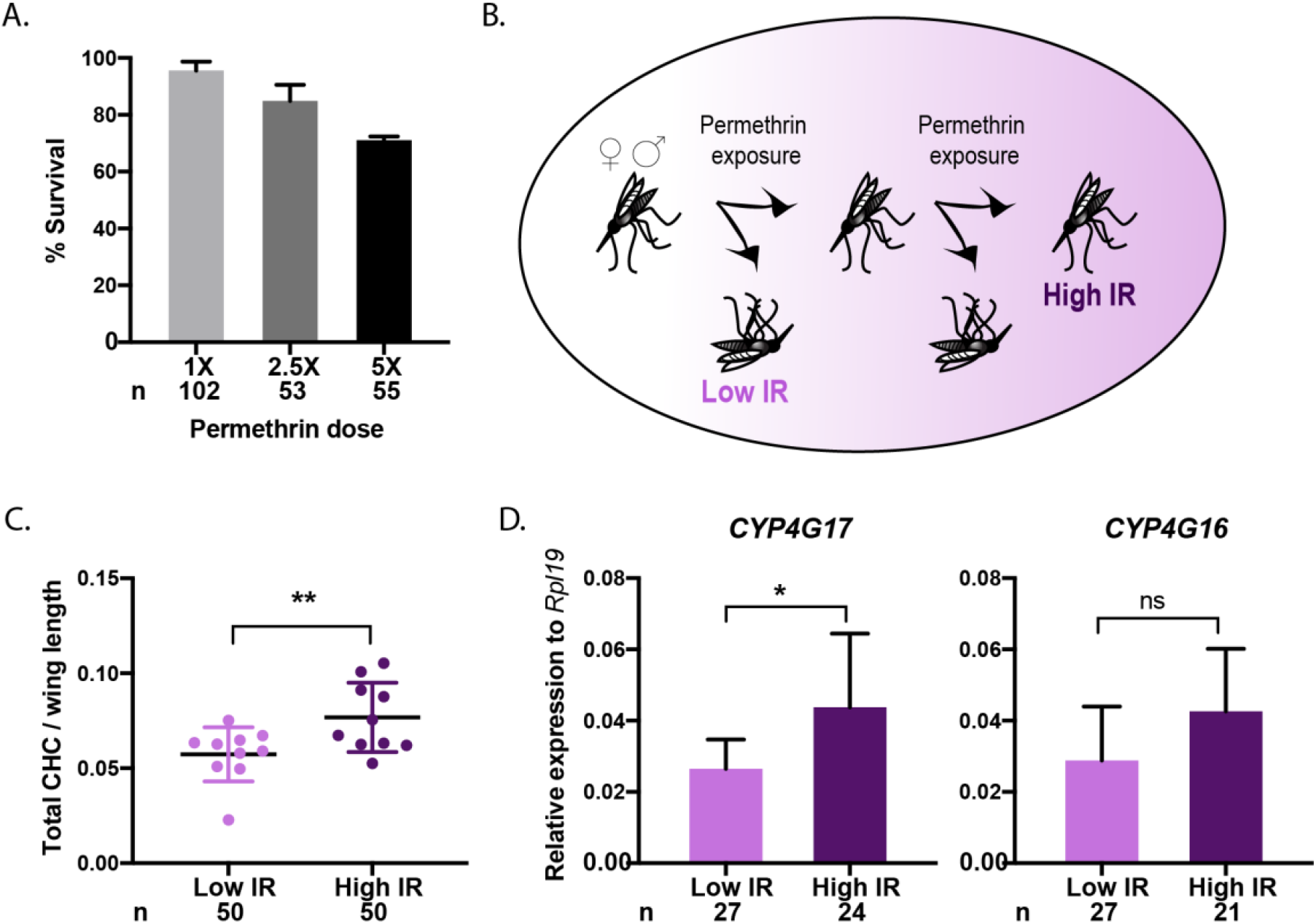
CHC levels are correlated with insecticide resistance intensity in field-derived *An. coluzzii*. (A) Insecticide resistance bioassays using 1X, 2.5X, or 5X permethrin-impregnated papers show high insecticide resistance in *An. coluzzii* adults collected as larvae from VK5 breeding sites. Scheme of experimental design showing how mosquitoes from larval collections were categorized as Low IR or High IR based on their survival to two sequential permethrin exposures. High IR mosquitoes show higher abundance of CHCs compared to Low IR mosquitoes (Generalized Linear Model, *p*=0.0083) after normalizing for wing length and accounting for sex within the model (*p*=0.2651). Mean and SD are shown. (D) High IR females also show higher transcript abundance of *CYP4G17* (unpaired t-test, *p*=0.0346), but not *CYP4G16* (unpaired t-test, *p*>0.05) by qRT-PCR. Bars represent mean and SD; *n* represents total number of mosquitoes.

We next tested whether CHC abundance is correlated with insecticide resistance intensity in the same population using age-matched *An. coluzzii* mosquitoes collected from larval breeding sites. We first separated mosquitoes according to their insecticide resistance intensity (Low or High insecticide resistance (IR)) based on their survival to two consecutive permethrin exposures. We classified as Low IR mosquitoes those who died after a single exposure to permethrin, while High IR mosquitoes were those who also survived a second exposure (**Figure 3B).** Permethrin doses used were different in males and females, as females survived longer to the same doses **(Supplementary Figure 3A)**. Data from males and females were pooled to increase sample size as we saw no difference in total CHC abundances between sexes **(Supplementary Figure 3B)**, though some qualitative differences were observed **(Supplementary Table 3)**. We detected a 1.33-fold increase in the total abundance of CHCs in High IR mosquitoes compared to the Low IR groups from the same population (Generalized Linear Model, *p*=0.0083) (**Figure 3C).** High IR mosquitoes were larger **(Supplementary Figure 3C)**, however neither wing length nor sex significantly explained variance in total CHC abundance when factored into a Generalized Linear Model as independent variables **(Supplementary Table 2C)**. Further, we accounted for size differences by normalizing CHCs to wing length **(Figure 3C)**.

To gain further evidence of cuticular resistance in those populations, we also investigated transcript levels for the cuticular resistance genes *CYP4G16* and *CYP4G17* among High and Low IR females. Sample collection and quality was not sufficient for data collection from males. *CYP4G17* expression was significantly higher in the High IR group (unpaired t-test, *p*=0.0346), while differences in *CYP4G16* expression trended in the same direction but were not significant **(Figure 3D)**. All together, these data provide evidence for cuticular resistance in this mosquito population, confirming published studies^28,33^, and show a direct association between CHC abundance and insecticide resistance intensity in age-matched mosquitoes.

## Discussion

In *Anopheles* mosquitoes, the traits defining male attractiveness and competitiveness during mating are not well understood. Here we show that males that are successful in mating swarms have higher abundances of CHCs, which could act as contact pheromones during interactions with females. Our data indicate that though relative abundances of different CHCs are similar among *An. coluzzii* males, greater total abundance is associated with mating success **(Figure 1B).** While CHC quantity could reflect increased male fitness involved in male-male competition, it is perhaps more plausible that this is a trait associated with female mate choice because CHCs are commonly used as contact pheromones in other insects^18^, and because of documented female rejection behavior during close range interactions^4,9,10^. This is supported by recent findings that show *An. stephensi* males treated with heptacosane, an abundant CHC also detected in our analyses, have higher insemination rates compared to control treated males^6^, suggesting that CHCs render males more attractive to females rather than reflecting increased male-male competitiveness (which is unlikely to be conveyed by exogenous addition of CHCs). *An. coluzzii* females may only assess the abundance of one or a few important compounds when evaluating male fitness, but there is precedent for association of total CHC abundance with mating outcomes. In *Drosophila serrata*, for example, overall CHC levels increase under conditions of sexual selection^36^, while in *Gnatocerus* flour beetles and *Cyphoderris* sagebrush crickets, total abundance is thought to play a role in mate choice, though CHC composition is also involved^36–38^. Notably, the upregulation of the CHC biosynthetic pathway has advantages beyond mating success. These compounds form a waxy seal on the exterior epicuticle that regulates permeability to water, insecticides, and other chemicals, and may therefore provide benefits in conditions of environmental stress. Indeed, in a region of Burkina Faso where high pressure from LLINs and indoor residual spraying have caused the emergence and spread of a number of insecticide resistance mechanisms, including cuticular thickening^27,28^, we found that males that were successful in mating were more resistant to pyrethroid insecticides **(Figure 2).** Consistent with previous studies, we also detected the upregulation of an important marker (*CYP4G17*) of cuticular resistance^27^ in mosquitoes that are more resistant to pyrethroids **(Figure 3D).** Combined, this evidence points to the possibility that CHC abundance may be not only be selected for by insecticide pressure, but also propagated by the increased mating success of individuals that possess cuticular resistance traits, unveiling an unexpected link between sexual selection and the failure of our best malaria control tools. Although most insecticide-based interventions are mainly targeted at adult females, adult males also rest indoors^39^ besides being exposed to insecticides during larval development, so increased CHC abundance is likely to be directly beneficial by increasing both the likelihood that males will survive long enough to mate and their chances of being successful during swarming events.

Our finding that CHCs act as dual traits in *An. coluzzii*, with roles in both mating behavior and in withstanding environmental pressures, is consistent with reports in *Drosophila* where hydrocarbons involved in mate choice are thought to have evolved differentially in different species based on environmental conditions like aridity of their ecological niches^19^. Similarly, different environments may explain why cuticular resistance has not been detected in all *Anopheles* populations, although it is also possible that the presence of this mechanism may be under-reported due to the lack of simple and reliable molecular diagnostic tools. Increased CHC production is likely to impose fitness costs that can only be offset in specific environmental conditions such as insecticide usage or aridity, and indeed there is evidence that CHCs also contribute to desiccation tolerance in *An. gambiae*^40–42^. Fitness costs of CHC production have been reported in *Drosophila*, where there appears to be a trade-off between CHC abundance and oogenesis, to the point that in the absence of sexual selection total CHC content is reduced in both male and female flies^36,43^. In the Vallée du Kou area where we conducted our studies, in addition to heavy insecticide usage mosquitoes are exposed to other significant environmental factors like a dry season that subjects mosquitoes to desiccation stress. Both these conditions are likely to promote increased CHC production even when a balance must be struck between abundance of these cuticular pheromones and their presumed fitness costs.

It is important to note that additional insecticide resistance mechanisms exist in this mosquito population including metabolic and target site resistance^33,34^. We cannot exclude that they contributed to the survival differences in our experiments, nor do we argue that cuticular resistance is necessarily equally or more dominant as a mechanism. However, the fact that we detected CHC differences between mosquitoes with low and high insecticide resistance **(Figure 3C)**, despite the potential contributions of other mechanisms, suggests cuticular resistance is important. In future studies, it will be critical to explore how different ecological factors shape mating success and sexual selection in *Anopheles* vectors from other malaria-endemic regions. Using wild-caught mosquitoes from natural swarms in our study has some inevitable limitations. We cannot be certain that males in our unmated group had not mated on previous nights, and we assume that mating does not change the CHC profile of males, an assumption supported by previous evidence^26^. Relatedly, due to the lack of appropriate high resolution age-grading technology, we cannot control for the age of males caught from natural swarms. Although we cannot rule out age as a confounding factor entirely, this parameter is not likely to critically influence our results due to the following arguments. Firstly, it has been shown that age is not associated with mating in natural swarms^15^. Moreover, the same study showed that 80-90% of *An. gambiae* swarming males are >4 days old and insemination rates are highest using males between 4 and 8 days of age^15^. Based on estimates of daily survival, as few as 5% of males may live longer than 8 days^44^, and we therefore expect that the vast majority of our cohort of swarming males are between 4 and 8 days old. Secondly, insecticide resistance does wane with age, but in the range of 4-8 days of age these effects are subtle, at least in females^45^. Lastly, though the impact of age on the total CHC abundance in males is not fully understood, a previous study observed an increase in CHCs with age in *An. gambiae* females^40^. Together these studies show that effects of age on CHC abundance and insecticide resistance may act in opposing directions, making it highly unlikely that age explains our findings here. Importantly, when age was controlled by using mosquitoes from larval collections from breeding sites in the same region, we still observed increased CHC abundance in mosquitoes that were more resistant to permethrin **(Figure 3B)**.

Finally, our findings that males with reduced CHC levels are less competitive in mating swarms also have repercussions for vector control strategies currently in the design stage that propose to release sterile or genetically modified males for malaria control. Given our results, laboratory-derived males lacking insecticide resistance mechanisms such as cuticular resistance would likely be less successful when mating in swarms alongside wild males, whether at the level of female mate choice or male competition. This reinforces the need to backcross laboratory mosquitoes sufficiently into the local genetic background before release, and to take into consideration the presence and mechanisms of insecticide resistance in the field population when planning the release of modified males.

## Materials and Methods

### Mating captures from natural swarms

Using small nets, mating couples of *An. coluzzii*, a subset of which were genotyped using primers from Santolamazza et al. 2008^46^, were manually caught from swarms in Vallée du Kou village 7 by trained personnel^47,48^. Samples collected for analysis of CHCs in swarming males (Figure 1) were obtained during September 2017, while samples collected in September 2018 were used to determine insecticide resistance intensities in swarming males (Figure 2). During collections, the nets were verified to contain one male and one female prior to being mouth-aspirated from the net into a small cup covered with netting. Nets that contained more than one male were discarded. For the unmated control groups, males were collected by one or several sweeps of a net through the swarm between 3 and 15 minutes into the swarm for the peak swarm time point, or 17-20 minutes into the swarm for the late swarm time point. Males from these sweeps were aspirated from nets into cups. Mosquitoes were given cotton soaked with 10% sugar solution and transported from field sites to an insectary in a vehicle. There they were additionally given cotton soaked in water overnight. CHCs were collected from these males 24 hours later.

### GC-M samples

For all GC-MS samples, pools of five mosquitoes were submerged in 200μL hexane for 30 min. Hexane was evaporated and samples were stored at room temperature. Just prior to GC-MS, samples were resuspended in 200μL hexane containing pentadecane as an internal standard of known quantity (1.53μg/sample). 3μL sample was injected into an Agilent fused-silica capillary column of cross-linked DB-5MS (30m × 0.25mm × 0.25μm). The GC conditions were as follows: inlet and transfer line temperatures, 290°C; oven temperature program, 50°C for 0.6 min, 50°C/min to 80°C for 2 min, 30°C/min to 120°C, 5°C/min to 310°C for 20 min, 50°C/min to 325°C for 10 min; inlet helium carrier gas flow rate, 1mL/min; split mode, splitless. These conditions are optimized for detection and resolution of lower chain length molecules.

### GC-MS analysis

A CHC accurate-mass target database was built based on the retention time comparison method and on the characterized ions reported by Caputo *et al.* 2005 to identify the compounds from the GC-MS run^49^. The relative response of the peak area of the extracted ion chromatogram of a target relative to the pentadecane internal standard was used to generate a quantitative value for each compound, called the response ratio.

### Absolute abundance

We compared this value between samples to look at the differences in absolute quantity of each CHC, after accounting for wing length of the mosquitoes in each sample. The sum of response ratios for all CHC compounds identified in a sample was used to compare total abundance of CHCs. We only included in our analyses compounds that were identified in at least 80% of all samples for each run. Data from one day of sample collection from swarms was excluded because all samples failed to detect 8 compounds that were detected in samples from all other collection days, and also expressed less than 25% of the total CHC abundance compared to the average. Statistics were performed as follows: For mated versus unmated comparisons, data were checked for normality prior to running One-Way ANOVA with Tukey’s multiple comparisons for mated versus peak unmated versus late unmated comparisons. These data were also incorporated into a Generalized Linear Model to determine whether there were significant differences between males captured from different swarms or on different nights (no significant effects were determined) as well as to account for wing length (prior to normalization). For Low IR versus High IR comparisons: data were checked for normality prior to running a Generalized Linear Model for Low IR versus High IR comparisons accounting for sex within the model. A Generalized Linear Model was also run to account for wing length prior to normalization, in addition to sex. For mosquitoes binned according to insecticide resistance intensity, animals in the Low IR groups tended to have fewer legs remaining post-exposure to permethrin, so the number of legs was normalized by removing the appropriate number of legs from High IR mosquitoes prior to hexane extraction to mirror the Low IR group.

### Proportional abundance

We calculated and compared the relative abundance of each compound in all samples to determine whether the proportional representation of each compound was different between groups. To compare abundance of individual compounds, data was not distributed normally for all compounds so Mann-Whitney tests were used to determine statistical differences between groups, and a Benjamini-Hochberg correction for multiple comparisons was subsequently performed with Q=0.2.

### Wing length measurement

Wings were imaged and measured from the proximal wing notch to the distal tip of the third cross vein using ImageJ^50,51^. All measurements were taken by the same person for consistency. The response ratio (relative to pentadecane standard) of total CHCs was divided by wing length (mm) to give normalized values. After testing to verify that the data falls into a normal distribution, wing lengths were compared using unpaired t-tests or One-Way ANOVA with Tukey’s multiple comparisons in GraphPad Prism 8.4.3 (GraphPad Software Inc. USA).

### Insecticide resistance bioassays

Permethrin-impregnated papers were prepared at 0.75% (1X), 1.875% (2.5X), or 3.75% (5X) permethrin (Sigma-Aldrich PESTANAL^®^ analytical standard) concentrations, weight/volume. Standard WHO bioassay tubes were used to expose mosquitoes to permethrin-impregnated papers for a given period. For standard WHO bioassays^32^, mosquitoes were exposed to permethrin for one hour, and 24h recovery time was allowed before assessing mortality. For non-standard time-to-death assays, mosquitoes were exposed for either 30- or 60-min periods and monitored for survival after a given recovery time, as shown in Figure 2A. To compare survival duration, log-rank tests were performed in JMP Pro 13 (SAS Corp. US). Proportional Hazards models were also performed in JMP to rule out replicate differences driving significance and also to account for the contributions of wing length to differences in survival. For non-standard assays binning mosquitoes as Low IR or High IR, mosquitoes were exposed to permethrin in two subsequent intervals as follows: for females, 60 min 5X permethrin followed by 120 min rest, followed by 30 min 5X permethrin and another 120 min rest (Supplementary Figure 3A, upper panel). For males: 30 min 2.5X permethrin exposure followed by 120 min rest, 30 min 2.5X permethrin, 120 min rest (Supplementary Figure 3A. lower panel.). Survival was assessed after each rest period. Pilot experiments were used to determine permethrin doses that yielded approximately equal numbers of High and Low IR mosquitoes.

### Transcriptional analysis of insecticide resistance genes

After insecticide resistance bioassays were performed and categorized females as Low IR or High IR, whole bodies were placed in RNA*later*™ (Thermo Fisher Scientific) in pools of three. A 2-hour post-exposure time point was used to reduce degradation of mosquitoes that had died during exposure, while allowing some recovery time for those that were only knocked down. After 24h at 4°C, excess RNA*later* was removed, and samples were frozen at −20°C. Samples were later homogenized in 600μL TRI reagent^®^ (Thermo Fisher Scientific). RNA was then extracted according to modified manufacturer instructions, including three washes of the RNA pellet with 75% ethanol. Samples were treated with TURBO™ DNase (Thermo Fisher Scientific) prior to quantification with a NanoDrop Spectrophotometer 2000c (Thermo Fisher Scientific). At this stage, we determined that male samples yielded poor quantity and quality RNA, while female samples were adequate for cDNA synthesis and qRT-PCR. Briefly, cDNA synthesis was performed using approximately 2μg RNA per sample in a reaction volume of 100μL. Published primer sequences for *CYP4G16* and *CYP4G17* were obtained as previously described^27^. qRT-PCR reactions were run on a StepOnePlus thermocycler using SYBR-Green Master Mix (Thermo Fisher Scientific), with 300nM primers and 1:3 dilutions of cDNA in a 15μL reaction. *RPL19* was used as a house-keeping gene to normalize Ct values obtained for each sample. Expression values were then compared using unpaired t-tests using GraphPad Prism after verifying that data was distributed normally.

### Mosquito larval collections and rearing

Larval breeding sites in Vallée du Kou village 5 were surveilled daily during September 2018, and larvae were collected between the L3 and pupal stage. Predators and competing mosquito species were removed, and mosquitoes were reared from these stages in natural spring water with TetraMin^®^ powder (Tetra) fed daily. Mosquitoes were sex-separated as pupae under a light microscope and given *ad libitum* access to cotton soaked with water and a 10% sugar solution throughout adulthood with a 12h light 12h dark cycle.

### Statistics and reproducibility

Detailed statistical methods are described within the relevant methods sections. CHC data was collected from swarming males on three separate nights to ensure reproducibility, and survival of swarming males to insecticide exposure was evaluated a total of six times. CHC data was collected from Low and High IR from only one of three experiments, but RNA was collected from females from all three replicates.

## Data availability

All pertinent data is available within the manuscript or upon request.

## Acknowledgments

We thank the villagers of VK5 and VK7 for their assistance in capturing mosquitoes from within swarms, and members of the Catteruccia laboratory for comments on the manuscript. Funding for this study was provided by a joint Howard Hughes Medical Institute and Bill and Melinda Gates Foundation Faculty Scholars Award to FC (Grant ID: OPP1158190), by the National institutes of Health (NIH) (award number: R01 AI104956) to FC and by a fellowship of the National Sciences and Engineering and Research Council of Canada to KA. The findings and conclusions within this publication are those of the authors and do not necessarily reflect positions or policies of the HHMI, the BMGF, the NIH or the NSERC.

