## Supplementary Figures and Tables for "Cuticular hydrocarbons are associated with mating success and insecticide resistance in malaria vectors"

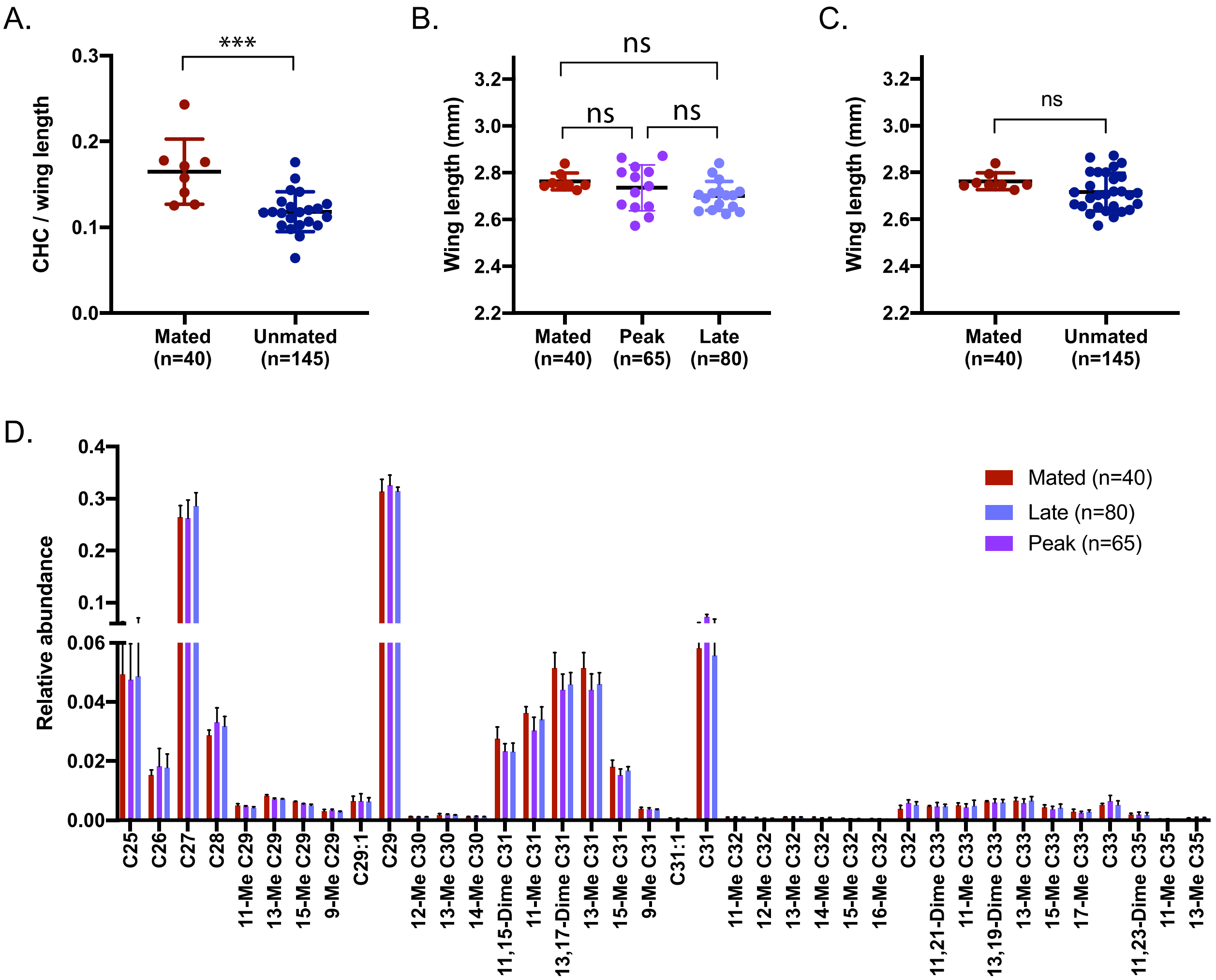


**Supplementary Figure 1**: **Neither body size nor proportional abundances of CHCs are different between mated and control groups.** (A) CHC abundance is higher in mated males compared to all unmated groups when peak and late time points are combined (Un-paired t-test, $p$=0.0003). Mean and SD are represented. (B) Wing lengths are not different between either late or peak unmated males compare to mated males (Tukey's multiple comparisons, $p$>0.05 for all comparisons). The mean wing length of each pool of 5 males is represented as each data point. Mean and SD are shown. (C) Wing lengths are not different between mated males and unmated males when peak and late time points are combined (Un-paired t-test, $p$>0.05). The mean wing length of each pool of 5 males is represented by each data point. Mean and SD are shown. (D) The proportional representation of all CHCs is not different between mated and unmated individuals from the peak or late time point of the swarm (Mann-Whitney tests, $p>0.05$, medians with interquartile ranges represented). *n* describes total number of mosquitoes.

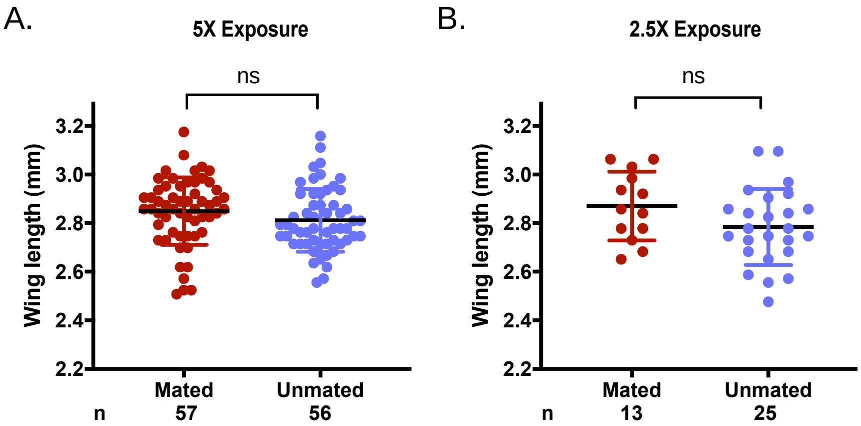


**Supplementary Figure 2: Wing lengths of mated and unmated males challenged with permethrin.** Wing lengths of individual males exposed to permethrin in (A) 5X continuous exposure experiment or (B) 2.5X intermittent exposure experiment do not show significant differences (unpaired t-tests, $p>0.05$, bars represent mean and SD). n represents total number of mosquitoes.

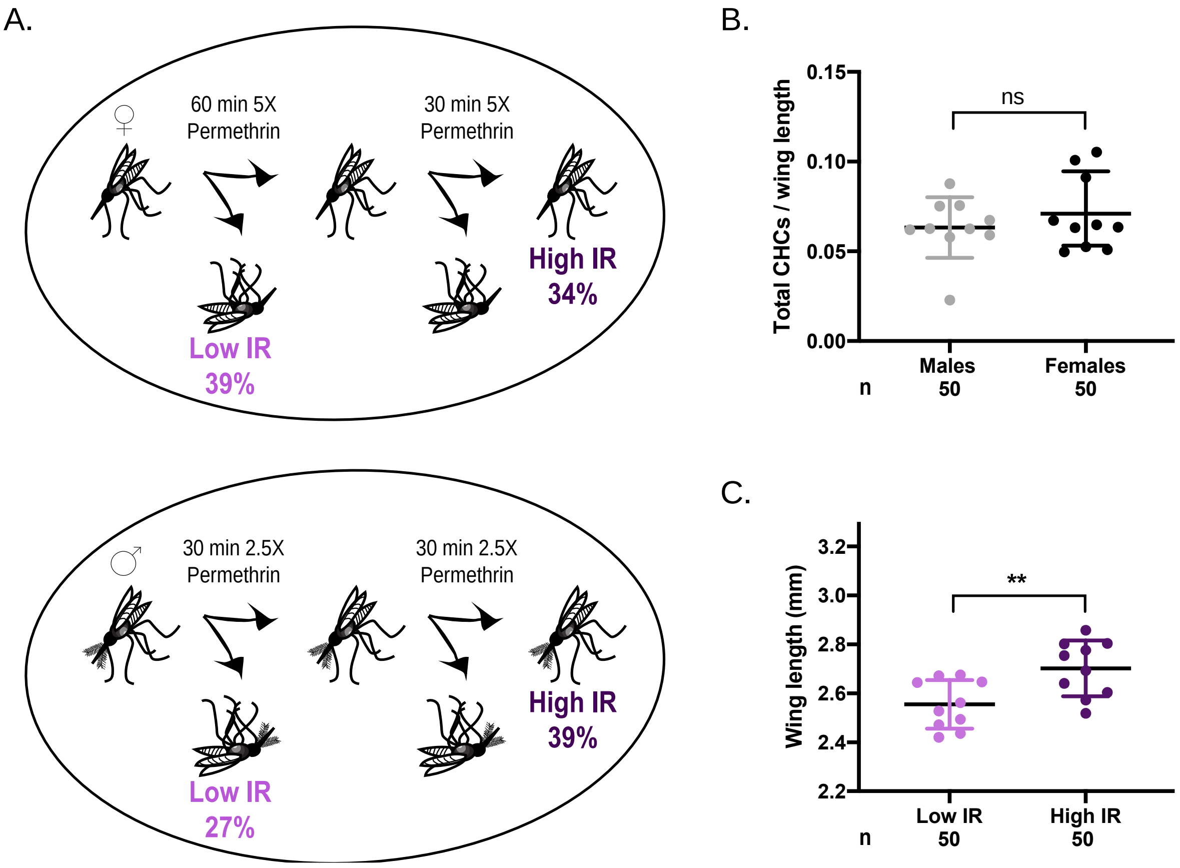


**Supplementary Figure 3: Wing lengths and CHC abundances of Low and High IR mosquitoes** (A) Schematics of procedure to separate females (i) and males (ii) into Low and High IR resistance groups based on their survival to two consecutive exposures to permethrin, with percentages showing the proportion of the population falling into either Low IR (died after the first exposure) or High IR (survived both exposures). (B) Total CHC abundances are the same in males compared to females, normalized to wing length (unpaired t-test, $p>0.05$, bars represent mean and SD. (C) Average wing lengths of each pool of 5 mosquitoes submitted for GC-MS are greater in High IR compared to Low IR groups (unpaired t-test, $p=0.0066$, bars represent mean and SD). *n* represents total number of mosquitoes.

**Supplementary Table 1**: **Absolute abundances of CHCs in mated and unmated *An. coluzzii* males from natural swarms.** Values show response ratios normalized to wing length. *p* values are calculated by Mann-Whitney tests, and then corrected for multiple comparisons using Benjamini-Hochberg procedure. Significant corrected *p* values (*p*<0.05) are shown in bold.

| | **Abundance** | | | ***P* value (Mann-Whitney)** | | ***P* value (Benjamini-Hochberg corrected)** | |
|---|---|---|---|---|---|---|---|
| Compound | Mated | Peak unmated | Late unmated | Mated vs Peak | Mated vs Late | Mated vs Peak | Mated vs Late |
| C25 | 0.008303 | 0.005654 | 0.005585 | 0.0535 | 0.1924 | 0.075 | 0.09210526 |
| C26 | 0.002572 | 0.002135 | 0.002292 | 0.2381 | 0.3196 | 0.09342105 | 0.09473684 |
| C27 | 0.041469 | 0.034294 | 0.030597 | 0.0446 | 0.0523 | 0.06578947 | 0.07105263 |
| C28 | 0.00488 | 0.003899 | 0.003631 | 0.0638 | 0.0192 | 0.07894737 | **0.04605263** |
| **11-Me C29** | 0.000791 | 0.000572 | 0.000509 | 0.0077 | 0.0017 | **0.02236842** | **0.00789474** |
| **13-Me C29** | 0.001233 | 0.000853 | 0.000791 | 0.0077 | 0.0013 | **0.025** | **0.00526316** |
| **15-Me C29** | 0.000884 | 0.000681 | 0.000589 | 0.0082 | 0.0045 | **0.03026316** | **0.01710526** |
| **9-Me C29** | 0.000492 | 0.000397 | 0.000317 | 0.0093 | 0.0126 | **0.03289474** | **0.03684211** |
| C29:1 | 0.001053 | 0.000759 | 0.00073 | 0.0465 | 0.0382 | 0.06973684 | 0.06315789 |
| **C29** | 0.047758 | 0.038032 | 0.03596 | 0.0159 | 0.0009 | **0.04210526** | **0.00263158** |
| **12-Me C30** | 0.000217 | 0.000137 | 0.000136 | 0.0012 | 0.0002 | **0.00394737** | **0.00131579** |
| **13-Me C30** | 0.000343 | 0.000225 | 0.000209 | 0.0168 | 0.0034 | **0.04342105** | **0.01184211** |
| 14-Me C30 | 0.000188 | 0.000146 | 0.000127 | 0.0446 | 0.0071 | 0.06447368 | **0.01973684** |
| 11-Me C31 | 0.004451 | 0.002746 | 0.002847 | 0.0246 | 0.0808 | 0.05131579 | 0.08289474 |
| **11,15-Dime C31** | 0.006308 | 0.003572 | 0.00399 | 0.0126 | 0.0087 | **0.03552632** | **0.03157895** |
| **13-Me C31** | 0.008805 | 0.005304 | 0.005263 | 0.0077 | 0.0036 | **0.02631579** | **0.01447368** |
| **13,17-Dime C31** | 0.003175 | 0.001739 | 0.001848 | 0.0077 | 0.0036 | **0.02368421** | **0.01315789** |
| **15-Me C31** | 0.000662 | 0.000461 | 0.000409 | 0.012 | 0.0057 | **0.03421053** | **0.01842105** |
| 9-Me C31 | 8.04E-05 | 6.06E-05 | 5.74E-05 | 0.0535 | 0.0126 | 0.07368421 | **0.03815789** |
| **C31:1** | 0.009543 | 0.008639 | 0.006976 | 0.0077 | 0.0071 | **0.02894737** | **0.02105263** |
| C31 | 0.000146 | 0.000108 | 0.000102 | 0.4137 | 0.023 | 0.09605263 | 0.05 |
| 11-Me C32 | 0.000114 | 6.53E-05 | 6.45E-05 | 0.1213 | 0.0284 | 0.08552632 | 0.05657895 |
| 12-Me C32 | 0.008805 | 0.005306 | 0.005264 | 0.1042 | 0.0629 | 0.08421053 | 0.07763158 |
| 13-Me C32 | 0.000165 | 0.000107 | 0.000106 | 0.0368 | 0.0382 | 0.05921053 | 0.06184211 |
| 14-Me C32 | 0.000135 | 8.15E-05 | 0.000089 | 0.0134 | 0.068 | **0.03947368** | 0.08026316 |
| 15-Me C32 | 0.000102 | 4.49E-05 | 4.93E-05 | 0.0246 | 0.0192 | 0.05263158 | **0.04473684** |
| **16-Me C32** | 7.42E-05 | 2.94E-05 | 2.31E-05 | 0.0045 | 0.0016 | **0.01578947** | **0.00657895** |
| C32 | 0.000551 | 0.000676 | 0.0006 | 0.6452 | 0.9761 | 0.09868421 | 0.1 |
| **11-Me C33** | 0.000776 | 0.000505 | 0.00051 | 0.0159 | 0.023 | **0.04078947** | **0.04868421** |
| 11,21-Dime C33 | 0.000851 | 0.000484 | 0.000549 | 0.1614 | 0.0382 | 0.08947368 | 0.06052632 |
| 13-Me C33 | 0.000948 | 0.000605 | 0.00066 | 0.0199 | 0.0702 | **0.04736842** | 0.08157895 |
| 13,19-Dime C33 | 0.001077 | 0.000629 | 0.000726 | 0.1257 | 0.0448 | 0.08815789 | 0.06710526 |
| 15-Me C33 | 0.000734 | 0.000403 | 0.000479 | 0.0246 | 0.0448 | 0.05394737 | 0.06842105 |
| 17-Me C33 | 0.000502 | 0.000237 | 0.0003 | 0.0077 | 0.0263 | **0.02763158** | 0.05526316 |
| C33 | 0.000939 | 0.000758 | 0.000616 | 0.5466 | 0.0523 | 0.09736842 | 0.07236842 |
| **11-Me C35** | 0.000314 | 0.000234 | 0.000222 | 0.0018 | 0.0028 | **0.00921053** | **0.01052632** |
| 11,23-Dime C35 | 4.56E-05 | 1.77E-05 | 2.19E-05 | 0.1614 | 0.1234 | 0.09078947 | 0.08684211 |
| 13-Me C35 | 0.000143 | 7.85E-05 | 8.75E-05 | 0.0288 | 0.0607 | 0.05789474 | 0.07631579 |

**Supplementary Table 2**: **Parametric model outputs.** (A) *P* values generated by Generalized Linear Model (Normal distribution). Y variable = CHC. Model effects include wing length and mating status (mated, or peak or late unmated). (B) Effect tests of proportional hazards. Y value = survival time. Model effects include wing length and mating status (mated vs unmated). (C) *P* values generated by Generalized Linear Model (Normal distribution). Y variable = CHC. Model effects include wing length, insecticide resistance status (Low IR vs High IR) and sex. In all models, interactions terms were removed as they were not significant ($p<0.05$).

| Model | Y value | Mating status | IR status | Wing length | Sex | Whole model | Df | Chi Square Value |
|---|---|---|---|---|---|---|---|---|
| A. Generalized Linear Model | **Total CHCs** | *p*=0.001 | - | *p*=0.0367 | - | *p*=0.0001 | 2 | 11.565 |
| B. Proportional Hazards | **Minutes survived** | *p*=0.1953 | - | *p*=0.6293 | - | *p*=0.3361 | 2 | 2.187 |
| C. Generalized Linear Model | **Total CHCs** | - | *p*=0.1031 | *p*=0.7476 | *p*=0.5244 | *p*=0.0105 | 3 | 11.2315 |

**Supplementary Table 3: Abundances of CHCs in Low IR or High IR mosquitoes.** Values show response ratios normalized to wing length.

| Compound | Abundance Low IR males ($x10^{-3}$) | Abundance High IR males ($x10^{-3}$) | Abundance Low IR females ($x10^{-3}$) | Abundance High IR females ($x10^{-3}$) |
|---|---|---|---|---|
| 13-Me C29 | 0.29624 | 0.40962 | 0.18132 | 0.25524 |
| 15-Me C29 | 0.22167 | 0.30402 | 0.16447 | 0.20917 |
| 11-Me C31 | 1.22651 | 1.54715 | 0.78088 | 1.3426 |
| 13-Me C31 | 1.49892 | 2.20471 | 0.79387 | 1.24947 |
| 15-Me C31 | 0.54675 | 0.78977 | 0.38437 | 0.58238 |
| 11-Me C33 | 0 | 0 | 0.30465 | 0.41216 |
| 13-Me C33 | 0.48426 | 0.42648 | 0.29724 | 0.38535 |
| 13,19-Dime C33 | 0 | 0 | 0.35104 | 0.33374 |
| 15-Me C33 | 0.28042 | 0.37498 | 0.15118 | 0.21748 |
| 13-Me C35 | 0 | 0 | 0.12137 | 0.17807 |
| C27:1 | 5.7665 | 7.06931 | 5.9586 | 8.27392 |
| C29:1 | 4.4635 | 5.49036 | 4.63286 | 6.84128 |
| C17 | 0 | 0 | 0.084602 | 0.085738 |
| C29 | 15.78802 | 19.82221 | 19.84549 | 27.57241 |
| C31 | 4.76514 | 5.85578 | 3.60508 | 4.62783 |
